# The chromatin modulating NSL complex regulates genes and pathways genetically linked to Parkinson’s disease

**DOI:** 10.1101/2023.01.16.523926

**Authors:** Amy R. Hicks, Regina Reynolds, Ben O’Callaghan, Sonia Garcia Ruiz, Ana Luisa Gil Martinez, Juan Botia, Helene Plun-Favreau, Mina Ryten

**Author notes:** These authors contributed to this study equally (Helene Plun-Favreau, Mina Ryten). Correspondence to: Mina Ryten, Full address: Department of Genetics and Genomic Medicine, Great Ormond Street Institute of Child Health, University College London, London, UK.

## Abstract

Genetic variants conferring risk for Parkinson’s disease have been highlighted through genome-wide association studies, yet exploration of their specific disease mechanisms is lacking. Two Parkinson’s disease candidate genes, *KAT8* and *KANSL1*, identified through genome-wide studies and a PINK1-mitophagy screen, encode part of the histone acetylating non-specific lethal complex. This complex localises to the nucleus, where it has a role in transcriptional activation, and to mitochondria, where it has been suggested to have a role in mitochondrial transcription. In this study, we sought to identify whether the non-specific lethal complex has potential regulatory relationships with other genes associated with Parkinson’s disease in human brain.

Correlation in the expression of non-specific lethal genes and Parkinson’s disease-associated genes was investigated in primary gene co-expression networks utilising publicly available transcriptomic data from multiple brain regions (provided by the Genotype-Tissue Expression Consortium and UK Brain Expression Consortium), whilst secondary networks were used to examine cell-type specificity. Reverse engineering of gene regulatory networks generated regulons of the complex, which were tested for heritability using stratified linkage disequilibrium score regression and then validated *in vitro* using the QuantiGene multiplex assay.

Significant clustering of non-specific lethal genes was revealed alongside Parkinson’s disease-associated genes in frontal cortex primary co-expression modules. Both primary and secondary co-expression modules containing these genes were enriched for mainly neuronal cell types. Regulons of the complex contained Parkinson’s disease-associated genes and were enriched for biological pathways genetically linked to disease. When examined in a neuroblastoma cell line, 41% of prioritised gene targets showed significant changes in mRNA expression following *KANSL1* or *KAT8* perturbation.

In conclusion, genes encoding the non-specific lethal complex are highly correlated with and regulate genes associated with Parkinson’s disease. Overall, these findings reveal a potentially wider role for this protein complex in regulating genes and pathways implicated in Parkinson’s disease.

## Introduction

An in-depth understanding of the genetic and pathophysiological mechanisms underlying neurodegenerative diseases is necessary to develop effective disease-modifying treatments. In the case of Parkinson’s disease, although 90-95% of cases are sporadic, historically much of the research into its genetic basis has focused on family-based linkage studies. Indeed, the identification of at least 23 genes with highly penetrant effects on Parkinson’s disease risk has succeeded in elucidating multiple biological pathways involved in its pathology. In particular, mitochondrial dysfunction and impaired protein degradation pathways are common themes. More recently, genome-wide association studies (GWASs) have identified 90 independent risk signals linked to Parkinson’s disease. Several of these had already appeared in familial studies, thereby highlighting important commonalities in the processes driving both types of the disease.^1^ However, a broader understanding of the molecular relationships between Parkinson’s disease loci is still lacking. Genes causally linked to the disease and involved in transcriptional regulation have the potential to provide such insights and shed light on key disease-relevant gene networks.

One such transcriptional regulator with strong links to Parkinson’s disease is *KAT8*. This gene was first linked to the disease through the identification of a risk signal on chromosome 16 (rs14235) with subsequent expression quantitative trait loci (eQTL) analysis suggesting the risk allele results in lower *KAT8* mRNA levels.^2,3^ Further GWAS analyses have again highlighted *KAT8* as a candidate gene, with recent colocalization and transcriptome-wide analyses strengthening the evidence for *KAT8’s* contribution to Parkinson’s disease.^1,4^ Importantly, KAT8 functions within two multiprotein complexes that regulate its activity and specificity, namely the male specific lethal (MSL) and non-specific lethal (NSL) complexes.^5^

Although the *KAT8-encoded* acetyl-transferase is thought to be the main catalytic driver in both complexes, differences in lysine specificity and in the genomic regions which are targeted can likely be attributed to subunits aside from KAT8 itself.^6^ This makes the other components of the MSL and NSL complexes of potential interest, with the latter particularly important in Parkinson’s disease as it contains KAT8 Regulatory NSL Complex Subunit 1 (KANSL1), another protein encoded by a Parkinson’s disease candidate gene.^3,7^ *KANSL1* is contained within the 970kb inversion polymorphism on chromosome 17q21, located within a linkage disequilibrium (LD) block of approximately 2Mb which gives rise to H1/H2 haplotype variation.^8^ The H1 haplotype has well established links to neurodegenerative disease, specifically progressive supranuclear palsy, Alzheimer’s and Parkinson’s disease.^9–11^ The precise mechanism underlying the link to Parkinson’s disease is disputed, with this risk frequently attributed to the adjacent tau-encoding *MAPT* as well as, more recently, a putative enhancer RNA expressed from within *KANSL1*.^12,13^ Moreover, the first GWAS of short tandem repeats in Parkinson’s disease found the strongest signal within *KANSL1*.^14^

Furthermore, both KAT8 and KANSL1 have been linked to mitophagy, the process by which defective mitochondria are identified and degraded and a key pathway implicated in Parkinson’s disease. Accumulation of mitophagy marker, phospho-ubiquitin (pUb, serine 65) has been detected in post-mortem diseased brains, whilst deficient mitophagy has been found in both sporadic and Mendelian patient-derived induced pluripotent stem cell (iPSC) models and even suggested to play a direct role in α-synuclein accumulation.^15–17^ Proteins involved in mitophagy, in particular PTEN-induced putative kinase 1 (PINK1) and parkin, are associated with early-onset autosomal recessive forms of the disease through mutations in their encoding genes, *PINK1* and *PRKN*.^18,19^ A biological screening assay of Parkinson’s disease GWAS candidate genes which measured PINK1-mediated mitophagy in neuroblastoma cells demonstrated significantly reduced pUb accumulation, parkin recruitment and phosphorylation, as well as lysosomal localisation of mitochondria following knockdown (KD) of both *KAT8* and *KANSL1*, thus demonstrating an important role of the NSL complex in mitochondrial quality control and Parkinson’s disease.^4,20^ However, there is some uncertainty regarding the precise molecular processes linking the NSL complex to PINK1-mediated mitophagy. There is evidence that components of the NSL complex can localise to mitochondria, though the most established function of the complex is in the nucleus, where it is involved in chromatin regulation.^21^ Thus, KAT8 and KANSL1 could operate to regulate the risk of Parkinson’s disease in multiple sub-cellular compartments.

In this study, we focused on the role of the NSL complex within the nucleus and tested our hypothesis that this complex operates as a master regulator of Parkinson’s disease risk. This idea is supported by existing evidence implicating KAT8-dependend lysine acetylation of primarily histone 4 in the regulation of a range of cellular processes, including DNA damage repair, and autophagy.^6,22–27^ To pursue this idea, we performed a series of *in silico* analyses which successfully predicted gene regulatory relationships between the chromatin modulating NSL complex and genes associated with Parkinson’s disease. These findings suggest a role for the NSL complex in modulating multiple pathological pathways and provide a useful framework for investigating potential gene regulatory mechanisms underlying disease risk associated with loci highlighted through GWASs.

## Materials and methods

### Gene selection

We collated three lists of genes, namely the NSL genes, genes causally associated with Mendelian forms of Parkinson’s disease and genes nominated through GWAS (Tab. 1). The nine genes encoding the NSL complex are widely published.^5^ An expert-curated list of genes linked to Mendelian forms of Parkinson’s disease and complex parkinsonism was obtained from PanelApp.^28^ Supplementary data published alongside the latest Parkinson’s disease GWAS was filtered for genes nominated by Mendelian Randomisation.^1^ Full gene lists are available here: https://github.com/amyrosehicks/NSL_PD_relationships (doi: 10.5281/zenodo.7525823).^29^

**Table 1.** Gene lists collated for frontal cortex gene co-expression network analysis. Genes missing from GCNs were not included in the search. Gene co-expression network (GCN), Genotype Tissue Expression (GTEx), non-specific lethal (NSL), United Kingdom Brain Expression Consortium (UKBEC).

| List | Source | Number of genes |  |  |
| --- | --- | --- | --- | --- |
|  |  | Total | Within GTEx GCN | Within UKBEC GCN |
| NSL complex | (Sheikh et al., 2019) | 9 | 9 | 8 |
| Mendelian Parkinson's disease | PanelApp: *Parkinson disease and complex Parkinsonism version 1.68* | 43 | 42 | 42 |
| Sporadic Parkinson's disease | Genes nominated by Mendelian Randomisation (Nalls et al., 2019) | 151 | 135 | 134 |

### Expression weighted cell-type enrichment analysis

Expression-weighted cell-type enrichment (EWCE) analysis was used to test whether NSL complex encoding genes are more highly expressed in specific brain-related cell types than would be expected by chance (https://github.com/NathanSkene/EWCE.git).^30^ Specificity values, representing the proportion of the total expression of a gene in one cell type compared to others, were calculated from two independent single nucleus RNA-sequencing datasets derived from human substantia nigra and medial temporal gyrus tissue, and were examined using the MarkerGenes github package.^31–34^ All data manipulation and visualisation was performed in R (v 4.0.5; RRID:SCR_001905) using the packages described here: https://github.com/amyrosehicks/NSL_PD_relationships.^29^

### Gene co-expression network analysis

We utilised two sources of public transcriptomic data for the generation of gene co-expression networks (GCNs), namely the Genotype Tissue Expression (GTEx, version 6) project (https://www.gtexportal.org/home/) and the United Kingdom Brain Expression Consortium (UKBEC, https://ukbec.wordpress.com/) data. GTEx contains samples originating from 13 CNS regions as well as other tissues and utilises Illumina sequencing, whilst UKBEC contains samples from 12 CNS regions assayed using Affymetrix arrays.^35,36^

Tissue-specific primary GCNs were built from these datasets using weighted gene coexpression network analysis (WGCNA), with k-means optimisation followed by functional and cellular specificity annotation.^34,37,38^ The completed GCNs formed part of the CoExp R package suite.^39^ Four secondary GCNs were also examined, for which the gene multifunctionality in secondary co-expression network analysis (GMSCA) R package was used to remove the contribution of neuronal, microglial, astrocytic and oligodendrocytic cell types from the expression matrix before GCN reconstruction.^40^

### Gene set enrichment analysis

For gene list enrichment analyses, we obtained p-values using Fisher’s Exact tests comparing the overlap in input genes and genes contained within each module, with false discovery rate (FDR) corrections for multiple testing.^39,40^ GCNs were annotated with gene ontology (GO) terms, categorized into biological process (BP), molecular function (MF) or cellular component (CC), within the CoExp R packages. GO terms were reduced to parent terms using the Rutils github package to aid visualization (doi: 10.5281/zenodo.6127446).^41^ Further analyses on gene lists derived from reverse engineering analysis were performed using the gProfiler2 R package (RRID:SCR_018190).^42^

### Reverse engineering gene regulatory network analysis

Using the Algorithm for the Reconstruction of Accurate Cellular Networks with adaptive partitioning (ARACNe-AP, RRID:SCR_002180), we inferred regulatory relationships between NSL complex genes and genes within specific modules of the primary GCNs.^43^ The original java package was adapted for use in R and executed with a p-value threshold of 1×10^-8^ and 10,000 bootstrap iterations. The strength of the association between regulators and regulons is denoted by mutual information values, which were compared using quantiles.^44^

### Stratified linkage disequilibrium score regression

Heritability is defined as the proportion of variation in a trait that can be attributed to inherited genetic factors.^45^ More specifically to this project, single nucleotide polymorphism (SNP)-based heritability refers to the variance that can be explained by any set of SNPs, such as those derived from a GWAS.^46^ We used stratified linkage disequilibrium score regression (LDSC, v.1.0.1., https://github.com/bulik/ldsc/wiki) to evaluate the enrichment of common SNP-based heritability for Parkinson’s disease across individual regulons derived from reverse engineering analysis and across regulon genes grouped according to cumulative frequency.^1,47,48^

Analysis was performed with parameters as described by Chen et al..^49^ Briefly, the baseline model of 97 annotations (v.2.2, GRCh37) to which annotations were added included only SNPs with minor allele frequencies over 5%. The major histocompatibility complex region was excluded due to its complicated LD patterns. Regression and LD reference panels utilised HapMap Project Phase 3 (https://www.sanger.ac.uk/data/hapmap-3/) and 1000 Genomes Project Phase 3 (https://www.internationalgenome.org/) European population SNPs respectively.^50,51^ Gene coordinates were extended by 100kb up and downstream of their transcription start and end sites to capture potentially relevant regulatory elements.^34^ The resultant regression coefficients (contribution of annotation after controlling for all other categories in the model) were used to calculate two-tailed p-values.

### Cell culture and siRNA treatment

We cultured wildtype (WT) SHSY5Y neuroblastoma cells (RRID:CVCL_0019) sourced from American Type Culture Collection (ATCC, RRID:SCR_00167) in Dulbecco’s Modified Eagle Medium (DMEM, Gibco, 11995-065) containing foetal bovine serum (10%, Gibco, 10500-064). For plating, cells were trypsinised, resuspended in culture media and counted using a Countess Automated Cell Counter.

Three types of siRNA were purchased as pre-designed siGENOME SMARTpools and transfected as per the manufacturer’s instructions: non-targeting/scrambled (D-001206-13), KAT8 (M-014800-00) and KANSL1 (M-031748-00). Each siRNA was diluted in FBS-free DMEM (final well concentration 50nM), mixed with DharmaFECT1 (Dharmacon, T-2001-03), 25μl added per well and incubated at room temperature for 30 minutes. Cell suspensions allowing for 12.5 x 10^3^ cells per well in a 96-well plate were prepared in culture media and 100μl added to each well on top of the siRNA mix. Following 72 hours incubation, media was removed and cells were lysed as per manufacturer’s instructions (see QuantiGene section below).

### QuantiGene multiplex assay

We used the QuantiGene multiplex system to simultaneously measure the expression of multiple genes in siRNA-treated SHSY5Y cell samples.^52,53^ Individual reagents and probe sets were purchased from ThermoFisher. Probes were directed against: i) five housekeeping genes used for normalisation, ii) two NSL complex genes used to quantify the KD, iii) all genes both causally linked to Mendelian Parkinson’s disease and complex parkinsonism, and contained within NSL regulons in the GTEx dataset, and iv) all genes nominated through GWAS which are expressed in brain and predicted to be regulated by at least three NSL complex genes (Tab. 2). *CTSB* was included as an exception, despite only appearing in two regulons, due to extensive literature implicating it in Parkinson’s disease.^54–56^ All GWAS-linked genes in this final list had mutual information values in the upper 50^th^ regulon quantile (Tab. 2).

**Table 2.** Details of QuantiGene 24-plex assay detecting changes in Parkinson’s disease-associated genes following NSL complex knockdown. All Mendelian disease-linked genes appearing in NSL regulons in the GTEx dataset were included, as well as GWAS genes expressed in brain and appearing in three or more regulons (*CTSB* was included as an exception due to extensive literature implicating it in Parkinson’s disease). Genotype Tissue Expression (GTEx), mutual information (MI), non-specific lethal (NSL), Parkinson’s disease (PD), United Kingdom Brain Expression Consortium (UKBEC).

| Symbol | Accession number | Type | Regulon frequency |  | Regulon MI quantile range |  |
| --- | --- | --- | --- | --- | --- | --- |
|  |  |  | GTEx | UKBEC | GTEx | UKBEC |
| *ATP5B* | NM_001686 | HK | - | - | - | - |
| *CANX* | NM_001746 | HK | - | - | - | - |
| *EIF4A2* | NM_001967 | HK | - | - | - | - |
| *HPRT1* | NM_000194 | HK | - | - | - | - |
| *PPIB* | NM_000942 | HK | - | - | - | - |
| *KANSL1* | NM_015443 | NSL | - | - | - | - |
| *KAT8* | NM_032188 | NSL | - | - | - | - |
| *PINK1* | NM_032409 | PD- mendelian | 2 |  | 0.544-0.648 |  |
| *ATN1* | NM_001940 | PD- mendelian | 1 | 1 | 0.942 | 0.966 |
| *ATXN2* | NM_002973 | PD- mendelian | 4 | 1 | 0.572-0.913 | 0.782 |
| *PARK7* | NM_007262 | PD- mendelian | 2 |  | 0.260 |  |
| *PLA2G6* | NM_003560 | PD- mendelian | 6 |  | 0.118-0.169 |  |
| *WDR45* | NM_007075 | PD- mendelian | 1 |  | 0.424 |  |
| *BIN3* | NM_018688 | PD- sporadic | 4 |  | 0.457-0.754 |  |
| *CCAR2* | NM_021174 | PD- sporadic | 3 |  | 0.729-0.839 |  |
| *CTSB* | NM_001908 | PD- sporadic | 2 |  | 0.825-0.899 |  |
| *DGKQ* | NM_001347 | PD- sporadic | 3 |  | 0.520-0.976 |  |
| *GAK* | NM_005255 | PD- sporadic | 3 |  | 0.864-0.977 |  |
| *NCKIPSD* | NM_016453 | PD- sporadic | 4 |  | 0.308-0.919 |  |
| *PGS1* | NM_024419 | PD- sporadic | 3 |  | 0.651-0.805 |  |
| *POLR2A* | NM_000937 | PD- sporadic | 3 | 1 | 0.463-0.533 | 0.617 |
| *QRICH1* | NM_017730 | PD- sporadic | 5 |  | 0.312-0.828 |  |
| *SETD1A* | NM_014712 | PD- sporadic | 4 | 2 | 0.294-0.869 | 0.0443-0.953 |
| *SH2B1* | NM_015503 | PD- sporadic | 5 |  | 0.385-0.978 |  |

Samples were prepared as per manufacturer’s instructions: working lysis mixture was prepared by diluting 1μl proteinase K per 100μl lysis mixture; cells were lysed by pipetting 150ul warm working lysis mixture per well (1:2 working lysis mixture to culture media); then plates were snap frozen on dry ice and stored at −80°C. The remainder of the assay was performed as indicated in the manufacturer’s protocol, with the exception that the Streptavidin R-Phycoerythrin conjugate (SAPE) binding step was completed at 51 °C.^53^ Plates were read using a Magpix (Luminex). This protocol is available on protocols.io (doi: dx.doi.org/10.17504/protocols.io.kqdg39ew7g25/v1).

Data analysis was performed by first subtracting the background, then normalising the signals obtained for the genes of interest to the geometric mean of the five housekeeping gene signals. Technical duplicates were included for each experimental repeat and outliers were identified and excluded using the ROUT method in GraphPad Prism (version 9, RRID:SCR_002798).^57^ Remaining duplicates were averaged and normalized to the SCR treated sample mean.

### Data availability

Raw data used to generate specificity matrices from substantia nigra and medial temporal gyrus are available at https://github.com/RHReynolds/MarkerGenes. Primary GCNs are available in the CoExpNets package (https://github.com/juanbot/CoExpNets) or on the CoExp website (www.rytenlab.com/coexp/Run/Catalog/). Details of the construction of secondary GCNs using the GMSCA package are available here: https://github.com/drlaguna/GMSCA. Stratified LDSC analysis utilised the LDSCforRyten package (https://github.com/RHReynolds/LDSCforRyten). Instructions for the use of ARACNe-AP were utilised from here: https://github.com/califano-lab/ARACNe-AP and the java package adapted for R is available alongside all code used for this paper here: https://github.com/amyrosehicks/NSL_PD_relationships.^29^

## Results

### Components of the NSL complex are highly expressed across all CNS regions and cell types

Using transcriptomic data provided by GTEx, the expression of genes encoding the NSL complex across the 13 CNS regions was analysed. This analysis demonstrated expression of all nine members of the NSL complex in all CNS regions, though we noted that *KAT8* and *OGT* were most highly expressed in the cerebellum (*median transcripts per million* (*TPM*) = 107.8 and 181.5) (Supplementary Fig. 1a). Differences in cell type-specific expression were also examined within two brain regions, namely the substantia nigra and medial temporal gyrus, utilising EWCE analysis and based on single-nuclei transcriptomic data.^31,32^ Consistent with expectation, the NSL complex genes displayed no overt differences in cell type-specific gene expression when considered collectively (Supplementary Fig. 1b).

### Components of the NSL complex cluster together in gene co-expression modules derived from human frontal cortex data

Given that there was no clear specificity of expression of NSL genes in CNS tissues or, more importantly, cell types, gene co-expression analysis was used to investigate the possibility of regional differences in co-expression that could explain selective neuronal vulnerability in Parkinson’s disease. This approach was based on the fact that genes with highly correlated expression tend to share biological relationships, and so GCN analysis can reveal otherwise hidden patterns in expression that reflect molecular and cellular processes.^39^ With this in mind, we used transcriptomic data generated by GTEx and covering 13 CNS regions to generate tissue-specific GCNs. Each GCN consisted of between 10-79 gene co-expression modules containing an average of 530 genes per module. Module membership values ranged from 0 to 1, with higher values indicating the expression of the gene is highly correlated with the module eigengene (ME).^58^

To identify gene co-expression modules of most interest, we investigated the enrichment of NSL genes within all 489 modules across all 13 GTEx CNS-relevant GCNs. We identified a significant enrichment of the NSL genes within the hippocampus ‘pink’ (*FDR corrected p-value* = 9.15×10^-5^, module membership range = 0.842-0.906) and frontal cortex ‘red’ modules (*FDR corrected p-value* = 5.86×10^-3^, module membership range = 0.775-0.904) (Fig. 1a, Supplementary Fig. 2). Importantly, the enrichment of NSL genes within the latter could be replicated in the corresponding UKBEC frontal cortex GCN within the ‘black’ module (*FDR corrected p-value* = 3.84×10^-4^, module membership range = 0.495-0.857) (Supplementary Fig. 3a). Given these findings, we focused on the GTEx and UKBEC frontal cortex GCNs and analysed the correlations between modules containing NSL genes. We found that several modules containing NSL genes, including the significantly enriched ‘red’ and ‘darkred’ (*FDR-corrected p-value* = 0.205, module memberships = 0.719 and 0.849) GTEx modules had highly correlated expression, as visualised in a ME dendrogram plot (Supplementary Fig. 4a,c). Again, this was replicated in the frontal cortex UKBEC GCN, where we noted high correlations in the expression of significantly enriched ‘black’ and ‘darkgreen’ (*FDR corrected p-value* = 0.270, module membership = 0.506) modules (Supplementary Fig. 4b,d). Taken together, these findings suggest that genes encoding the NSL complex are significantly co-expressed in the frontal cortex, a finding which is consistent with the importance of this brain region in Parkinson’s disease progression and dementia.^59–62^

**Figure 1.**
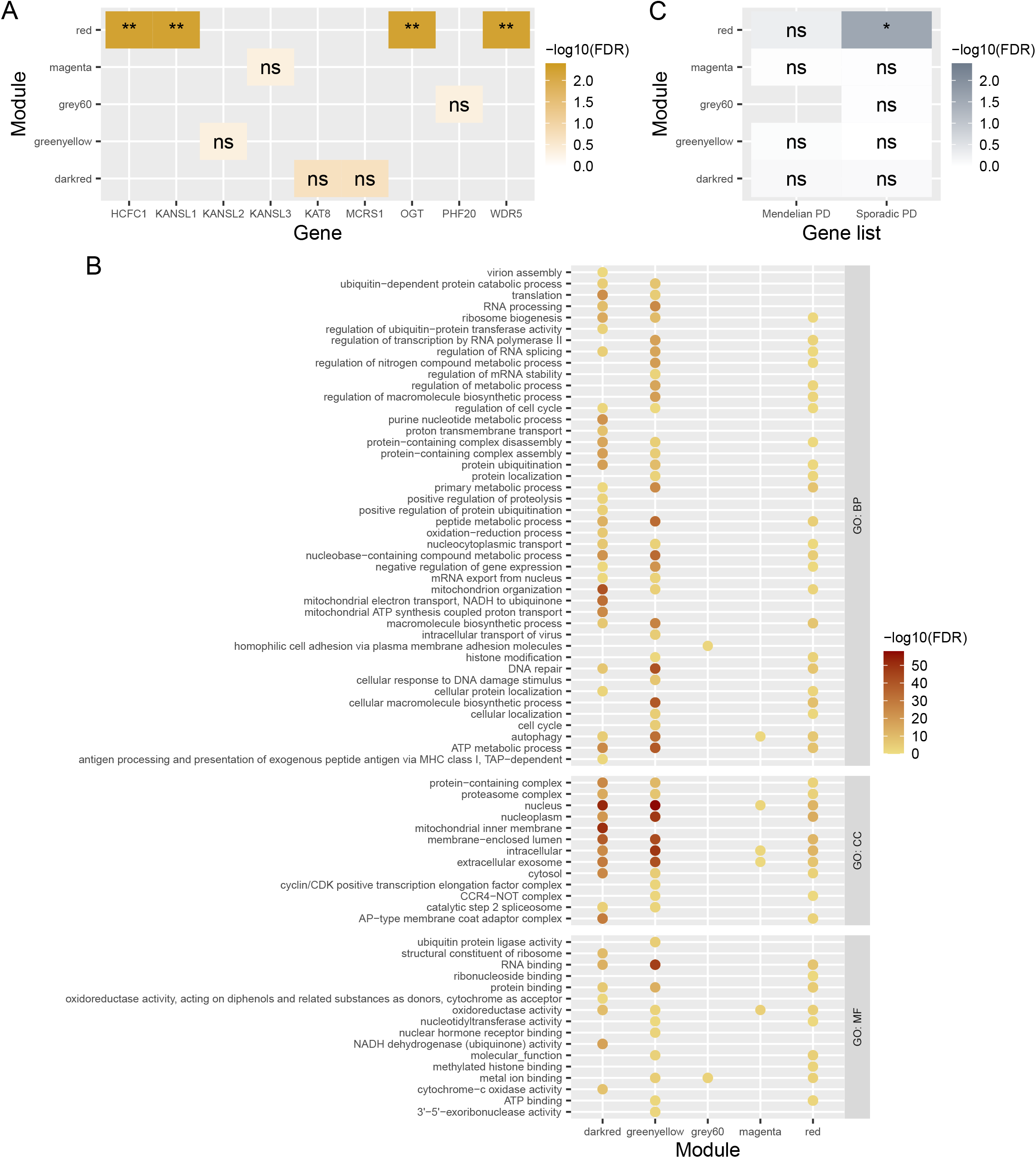
Exploration of GTEx frontal cortex gene co-expression network. (A) NSL gene enrichment analysis across GCN modules constructed using GTEx datasets. (B) GO BP, MF and CC term enrichments for GCN modules containing NSL genes. Each term has been uniformly reduced to a parent term in order to group together similar terms. Colour corresponds to the p-value of the most significantly enriched child term within the parent term. (C) Parkinson’s disease-associated gene set enrichment analysis, filtered for GCN modules containing NSL genes. Fishers exact test, displayed over a log scale (FDR-corrected p-values). ns, FDR > 0.05; *, FDR < 0.05; **, FDR < 0.01; ***, FDR < 0.001; ****, FDR < 0.0001. Biological process (BP), cellular compartment (CC), false discovery rate (FDR), gene co-expression network (GCN), gene ontology (GO), Genotype Tissue Expression (GTEx), molecular function (MF), non-specific lethal (NSL), Parkinson’s disease (PD).

### Co-expression analysis supports a role for the NSL complex in the regulation of both chromatin and mitochondrial function in human frontal cortex

Next, we focused on co-expression modules containing NSL complex genes to better understand their function in human brain. This was achieved by examining GO term enrichment within all five modules of interest in the GTEx frontal cortex GCN, but with the primary focus being the ‘red’ module. In total, these five modules were significantly enriched for 2899 GO terms (*FDR* < 0.05). Following term reduction, we noted enrichments of terms representing both the well-characterised nuclear-based and lesser-known mitochondrial-based role of the NSL complex. Consistent with expectation, nucleus-related terms, such as transcription coactivator activity (*FDR range* = 1.85×10^-5^-0.0227), were identified as enriched within the ‘red’ module, whilst the ‘darkred’ module was enriched for a range of mitochondria-related terms, including cytochrome-c oxidase activity (*FDR range* = 2.03×10^-8^-0.0128) and mitochondrial inner membrane (*FDR range* = 1.33×10^-51^-2.46×10^-3^) (Fig. 1b). The UKBEC GCN was also significantly enriched for several nuclear terms in the ‘black’ module, but lacked any mitochondrial terms (Supplementary Fig. 3b). Together these results indicated that both the chromatin- and mitochondria-related functions of the NSL complex were captured in these GCNs, indicating that these functions are likely to be active within human frontal cortex.

### Components of the NSL complex cluster together with Parkinson’s disease-associated genes in co-expression modules in human brain

Next, we explored the possibility that NSL genes are functionally related to genes associated with Parkinson’s disease. Parkinson’s-related genes were divided into those causally linked to Mendelian forms of the disease and those associated with sporadic Parkinson’s disease through the identification of GWAS risk loci in close proximity. The GTEx GCN was then tested for enrichment of the two disease lists. Gene set enrichment analysis revealed a significant enrichment of GWAS genes in the ‘red’ GTEx GCN module (*FDR* = 0.0269, module membership range = 0.553-0.907) (Fig. 1c). These results point to the possibility that important gene regulatory links exist between the NSL complex and genes associated with sporadic Parkinson’s disease.

### NSL complex activity in Parkinson’s disease may be most important in neuronal cell types

Although the genes encoding the NSL complex are expressed ubiquitously across different cell types, their gene regulatory relationships may nonetheless be cell type-specific, and only in specific cell types might there be a relationship with Parkinson’s disease genes. We used the GMSCA tool to test this possibility, and generated secondary GCNs, from which the contribution of four major cell types (neurons, astrocytes, microglia and oligodendrocytes) was removed in turn prior to network construction.^40^ As might be expected, this correction altered the clustering of NSL genes and their co-expression patterns with Parkinson’s disease-associated genes across secondary GCN modules (Fig. 2a,c). The sporadic Parkinson’s disease gene list was significantly enriched alongside *KANSL1* – as in the primary GCN – within the oligodendrocyte-corrected ‘green’ module (*FDR-corrected p-values* = 4.81×10^-3^ and 5.87×10^-3^ respectively) alone, whilst each of the neuron-, microglia- and astrocyte-corrected networks disrupted the relationship between NSL and disease genes (Fig 2a,c). This suggested that the relationship between these genes is likely to be active in the neurons, microglia and/or astrocytes.

**Figure 2.**
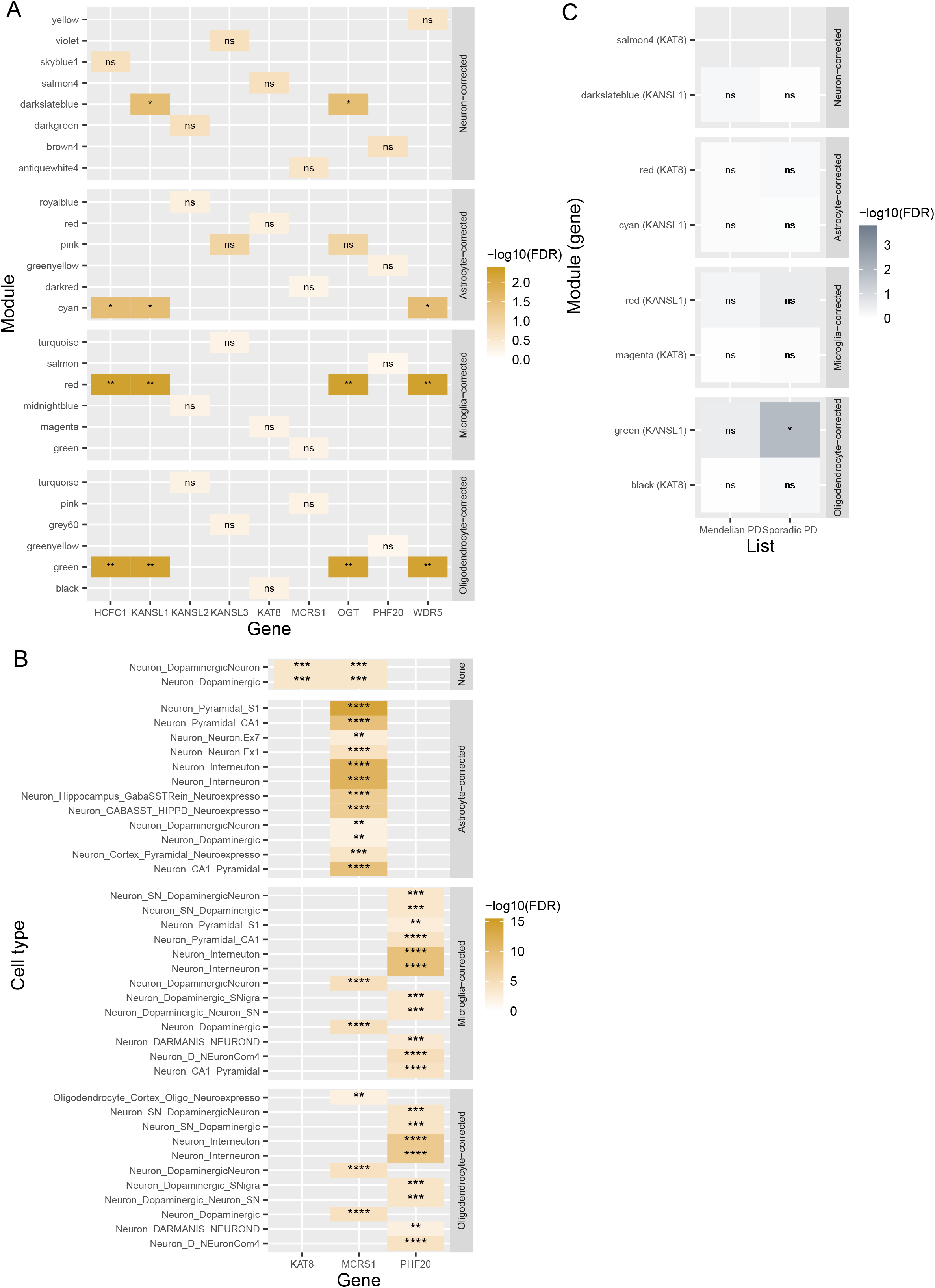
Exploration of GTEx frontal cortex secondary gene co-expression networks. (A) NSL gene enrichment analysis of secondary GCN modules constructed using GTEx datasets. (B) Cell type enrichment analysis of primary (labelled ‘none’) and secondary (labelled ‘corrected’) GCN modules containing NSL genes. NSL genes found within modules devoid of any cell type enrichment were not included. (C) Mendelian and sporadic Parkinson’s disease-associated gene enrichment analysis of secondary GCN modules, filtered for those containing *KAT8* and *KANSL1* (gene in brackets, no box indicates no genes present). Fishers exact test, displayed over a log scale (FDR-corrected p-values). ns, FDR > 0.05; *, FDR < 0.05; **, FDR < 0.01; ***, FDR < 0.001; ****, FDR < 0.0001. False discovery rate (FDR), gene coexpression network (GCN), Genotype Tissue Expression (GTEx), non-specific lethal (NSL), Parkinson’s disease (PD).

To identify which of the three cell types was most important for NSL-Parkinson’s disease coexpression, we examined the enrichment of cell type markers across all primary and secondary GCN modules. The ‘darkred’ module containing *KAT8* and *MCRS1* was enriched for markers of dopaminergic neuronal signalling (*p-value* = 3.23×10^-5^) in the GTEx primary GCN (Fig. 2b). *MCRS1* remained predominantly in modules enriched for different neuronal markers following the correction of microglial, astrocytic and oligodendrocytic signatures (*p-value range* = 3.92×10^-16^-7.42×10^-3^, module membership range = 0.8452-0.8529) (Fig. 2b). The primary GCN ‘grey60’ module containing *PHF20* lacked any cell type enrichment, contrasting to the secondary GCN ‘salmon’ and ‘greenyellow’ modules which were both enriched for multiple neuronal cell types following the correction of microglial and oligodendrocytic signatures respectively (*p-value range* = 3.99×10^-11^-5.11×10^-3^, module memberships = 0.8544 and 0.8539)(Fig. 2b). Although *KANSL1*-containing modules had no cell type enrichments, these results suggest the gene regulatory links between the NSL complex and genes associated with Parkinson’s disease may be most important in neuronal cell types.

### *In silico* analysis predicts the regulation of Parkinson’s disease-associated genes by members of the NSL complex

The genetic interactions modelled in GCNs are typically undirected in that causality is unassigned.^38^ However, it is already known that the NSL complex is highly important in the regulation of gene expression, suggesting that at least a proportion of the genes co-expressed with the NSL complex are regulated by it.^5^ We formally tested this possibility in silico with the tool Algorithm for the Reconstruction of Accurate Cellular Networks with adaptive partitioning (ARACNe-AP), which uses expression data to reverse engineer gene regulatory networks.^43,63^ By applying ARACNe-AP, we predicted the genes most likely to be regulated by the NSL complex amongst those contained within the ‘red’ and ‘darkred’ GTEx GCN modules (Supplementary Table 1).^43^ The resulting target gene lists, termed regulons, produced by this analysis ranged from 491 genes predicted to be regulated by *KANSL1*, to 1788 predicted to be regulated by *WDR5* in the GTEx dataset. As expected for genes encoding a protein complex which regulates gene expression, regulons showed significant overlaps with each other. The regulons of the four NSL genes contained within the ‘red’ GTEx GCN module all significantly overlapped (*FDR range* = 3.53×10^-67^-3.76×10^-4^), whilst the regulon of *KAT8* significantly overlapped only with that of *WDR5* (*FDR* = 2.98×10^-11^) (Fig. 3a).

**Figure 3.**
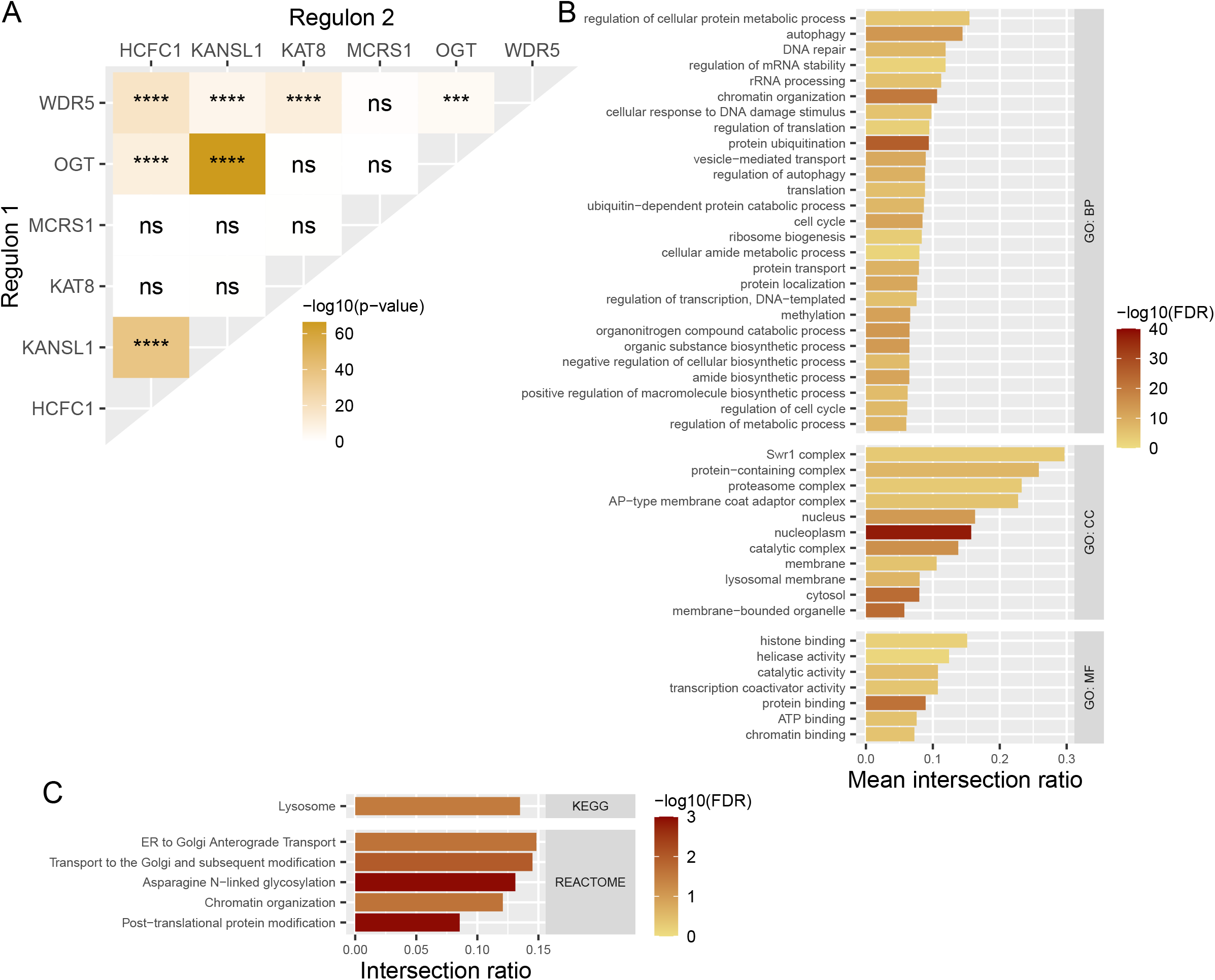
Characterisation of NSL complex regulons, derived from ARACNe-AP analysis of ‘red’ and ‘darkred’ GTEx primary gene co-expression network modules. (A) Overlaps of regulons of each NSL gene. (B) GO BP, MF and CC term enrichments for genes appearing in three or more NSL regulons. Each term has been uniformly reduced to a parent term in order to group together similar terms. Colour corresponds to the p-value of the most significantly enriched child term within the parent term and the x-axis denotes the mean ratio of genes intersecting with the term to total genes within each term. (C) REACTOME and KEGG term enrichments for genes appearing in three or more NSL regulons, filtered for those genetically linked to Parkinson’s disease. The x-axis denotes the ratio of genes intersecting with the term to total genes within each term. Fishers exact tests, displayed over a log scale (gene set enrichment p-values are FDR corrected). ns, FDR > 0.05; *, FDR < 0.05; **, FDR < 0.01; ***, FDR < 0.001; ****, FDR < 0.0001. Biological process (BP), cellular compartment (CC), false discovery rate (FDR), gene co-expression network (GCN), gene ontology (GO), Genotype Tissue Expression (GTEx), molecular function (MF), non-specific lethal (NSL), Parkinson’s disease (PD).

To reduce the impact of noise and focus on genes most representative of NSL complex activity, genes appearing in three or more regulons were collated and termed the NSL regulon (n = 1101). We then assessed this gene set for its role in Parkinson’s disease causation. Firstly, we noted that two Mendelian disease-associated genes (*ATXN2* and *PLA2G6*) and 12 sporadic disease-associated genes (*BIN3, CCAR2, DGKQ, GAK, GBAP1, IGSF9B, NCKIPSD, PGS1, POLR2A, QRICH1, SETD1A, SH2B1*) were contained within this dataset, though no significant enrichment was observed (*p-value* = 0.843 and = 0.619 respectively). Furthermore, we used stratified LDSC to assess the enrichment of Parkinson’s disease SNP-based heritability amongst all genes in the NSL regulon.^1^ Despite the relatively small gene set, we found that the regression coefficient – a stringent measure of the contribution to SNP-based heritability which accounts for underlying contributions of genetic architecture captured within the baseline model – had a p-value that fell just outside significance (*p-value* = 0.0519, Supplementary Tab. 2). This hinted at a link between Parkinson’s disease heritability and NSL complex activity. Individual regulons and gene lists divided according to cumulative regulon frequency were also tested, revealing a nominally significant enrichment of heritability within the *HCFC1* regulon (*p-value* = 0.0239, *FDR* = 0.143) (Supplementary Tab. 3).

Next, we assessed the NSL regulon for the enrichment of gene pathways of potential relevance to Parkinson’s disease. Using gene set enrichment analysis utilising parent terms to reduce redundancy, we found that genes contained within the NSL regulon were enriched for a range of terms including: autophagy (GO:0006914, *child term FDR range* = 3.11×10^-12^-0.0354), regulation of autophagy (GO:0010506, *child term FDR range* = 2.90×10^-4^-0.0123), and chromatin organisation (GO:0006325, *child term FDR range =* 7.50×10^-9^-0.0483) (Fig. 3b). Gene set enrichment analysis was also completed on individual regulons, revealing many of the same parent terms to be enriched across different regulons (Supplementary figure 5). To assess the association of Parkinson’s disease-relevant pathways with the regulons more directly, we specifically asked whether our enriched gene sets overlapped with a list of 46 gene sets genetically implicated in the disease through common genetic variation.^64^ This highlighted the enrichment of five disease-linked terms, including KEGG:04142 Lysosome (*FDR* = 0.0321) and REAC:R-HSA-4839726 chromatin organization (*FDR* = 0.0242), suggesting that the NSL regulon in frontal cortex is enriched for genes causally implicated in Parkinson’s disease (Fig. 3c).

### *In vitro* analysis confirms the regulation of Parkinson’s disease-associated genes by the NSL complex

Given the success of our in silico analyses, we wanted to validate some of the regulatory relationships identified *in vitro*, in particular focusing on genes causally associated with Parkinson’s disease and contained within the NSL regulon (Table 2). With this in mind, we measured the expression of 17 genes of interest in response to *KANSL1* and *KAT8* siRNA KD in a SHSY5Y cell line, a regularly used cell model for Parkinson’s disease research (Fig. 4, Supplementary Fig. 6).^4^ After confirming that *KANSL1* and *KAT8* KDs did not affect housekeeping gene expression levels, we found that both KDs significantly reduced *PINK1* expression (*p-value* < 1×10^-4^ and 0.0151), consistent with previous reports (Supplementary Fig. 6a,b).^4^ *BIN3, CTSB, DGKQ, NCKIPSD* and *PGS1* were also significantly reduced following *KANSL1* KD alone (*p-value range* < 1×10^-4^-0.0295), with reductions in *DGKQ* and *NCKIPSD* following *KAT8* KD also reaching significance (*p-value* = 4.55×10^-3^ and 4.25×10^-3^). Interestingly, *WDR45* expression followed an inverse pattern, with *KAT8* KD resulting in a significantly increase in *expression(p-value* = 3.31×10^-3^) (Fig. 4). Thus, seven of the 17 genes (41.2%) predicted to be regulated by the NSL complex using *in silico* analyses were indeed found to show significant changes in expression when *KAT8* or *KANSL1* expression was suppressed (Fig. 5).

**Figure 4.**
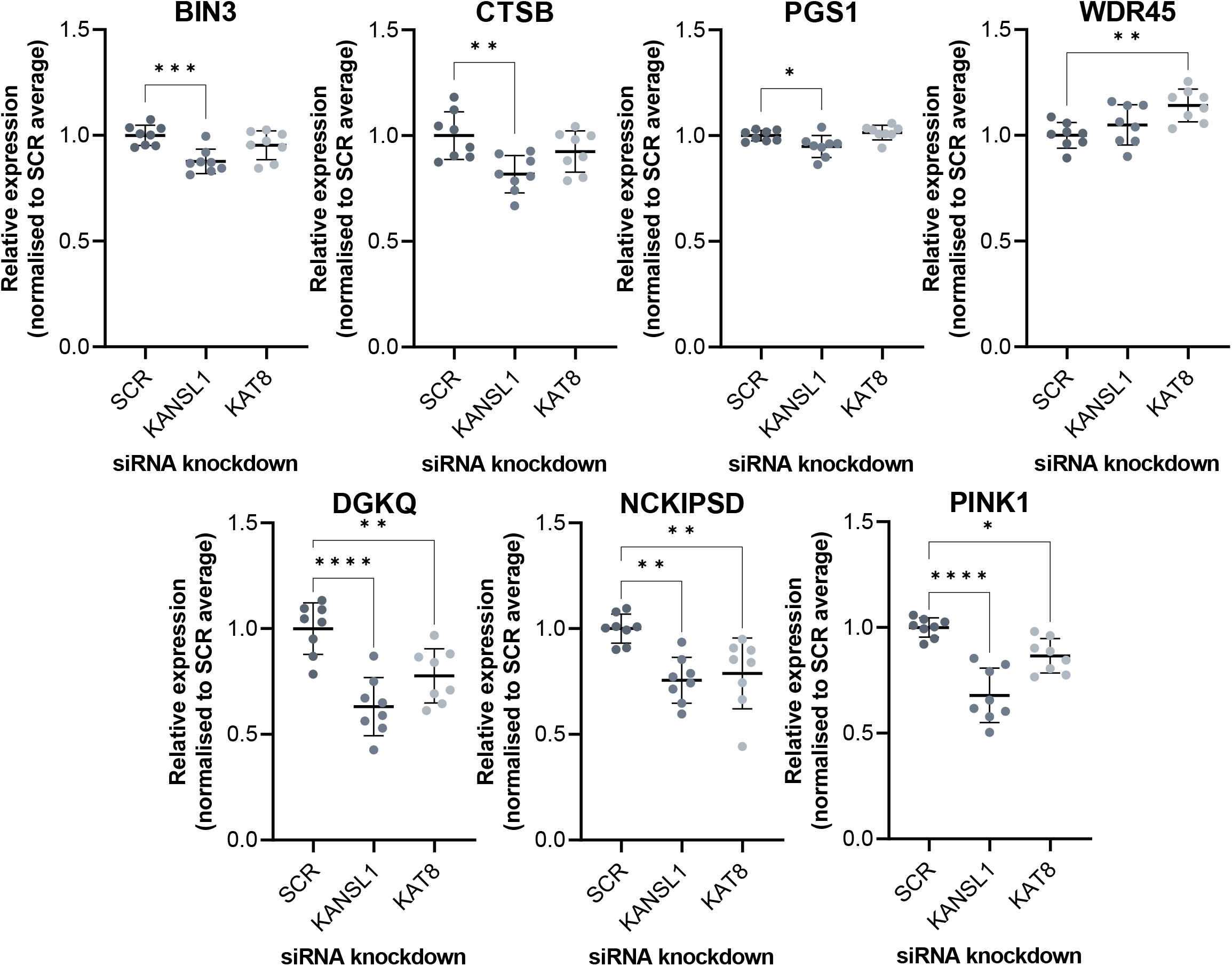
Changes in mRNA levels of Parkinson’s disease-associated genes following suppression of NSL complex genes. Measured using QuantiGene multiplex assay, normalised to average SCR control. One-way ANOVA with Dunnett correction for multiple comparisons, n = 8. *, FDR < 0.05; **, FDR < 0.01; ***, FDR < 0.001; ****, FDR < 0.0001. Non-specific lethal (NSL), scrambled (SCR).

**Figure 5.**
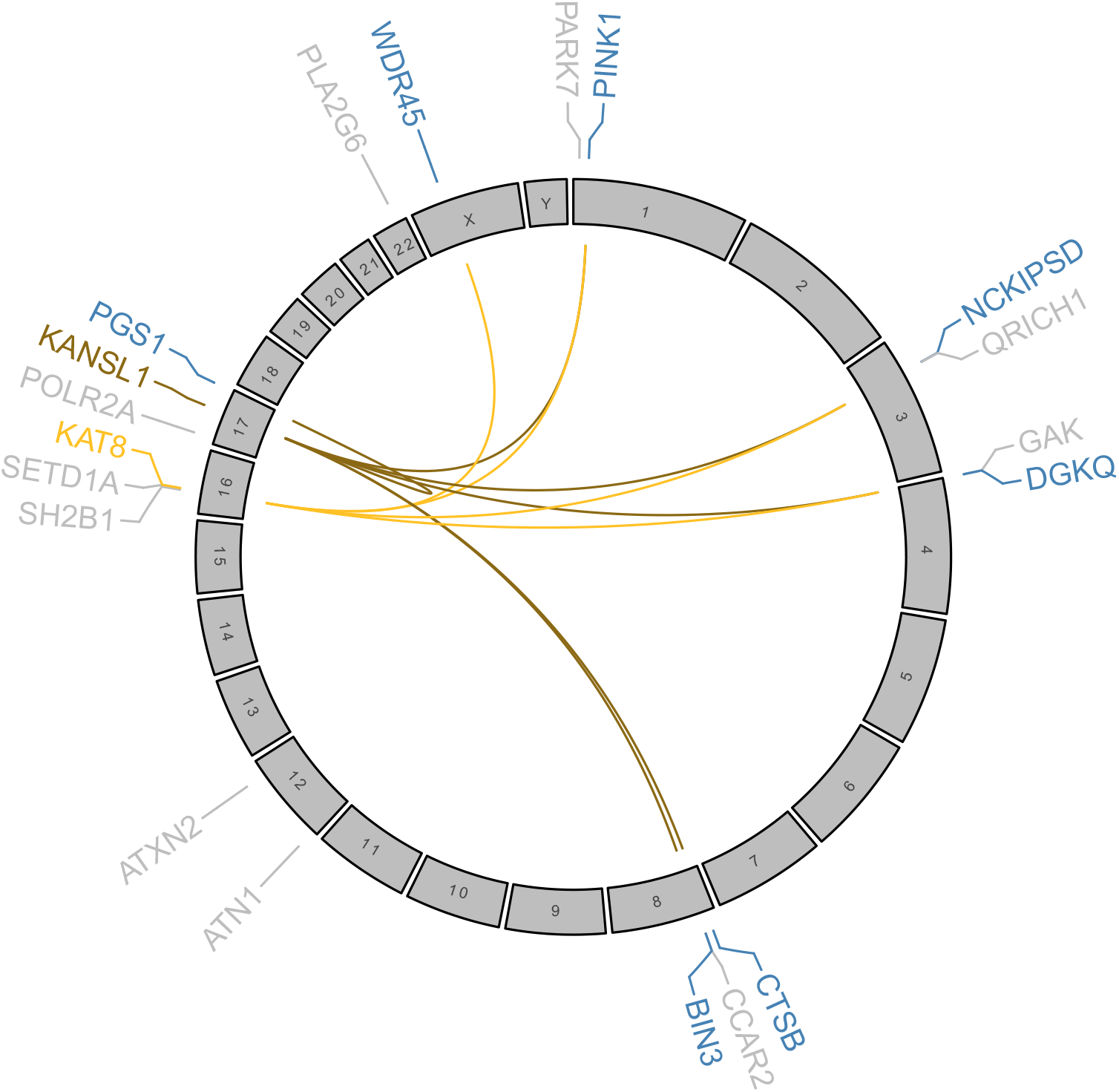
Regulatory relationship detected between genes encoding the NSL complex and genes associated with Parkinson’s disease. Genes highlighted were tested using QuantiGene multiplex assay. Colours correspond to classification of gene (blue, genes linked to Parkinson’s disease with NSL regulation detected; grey, genes linked to Parkinson’s disease without NSL regulation detected; lightest yellow, *KAT8* regulation; darkest yellow, *KANSL1* regulation). Non-specific lethal (NSL).

## Discussion

This project utilised publicly available transcriptomic data from human brain tissue to characterise the expression patterns of genes encoding the NSL complex and their relationships to genes genetically linked to Parkinson’s disease. First, NSL genes were found to cluster together with Parkinson’s-associated genes in GCN modules annotated for both chromatin and mitochondria-related functions. Second, these co-expression relationships appeared to be most associated with neuronal cell types. Third, a number of Parkinson’s-associated genes predicted to be directly regulated by multiple components of the NSL complex were subsequently validated in a relevant cell model.

Despite no clear specificity of expression being detected across tissues, the NSL genes showed specific co-expression patterns across GCNs produced from different brain regions. These results are consistent with the theory of the NSL complex having context-specific gene regulatory functions. Within the frontal cortex GCN, NSL genes clustered within GCN modules which were significantly enriched for nuclear GO terms, as expected for a chromatin regulating protein complex. The enrichment of mitochondrial GO terms within modules containing NSL genes adds supporting evidence for the less well-characterised role of the NSL complex based at mitochondria.^4,21^ Further, the enrichment of genes associated with sporadic Parkinson’s disease alongside NSL genes highlights a high level of correlation in gene expression. These results in particular support a functional interaction with the chromatin regulating NSL complex and point to a biological link between multiple genes highlighted through GWASs. It would be difficult to investigate such interactions in any way other than using post-mortem tissue given that this method captures the full background of the disease. However, this also means the results depend heavily on the numbers of samples available as this may limit the power of analyses. Particularly in the context of Parkinson’s disease, it may have been most interesting to investigate gene regulatory interactions within the substantia nigra, where much of the pathophysiological features of the disease are observed^65^. However, within the GTEx dataset studied here, there are only 63 substantia nigra samples compared to 108 frontal cortex.^36^

Secondary GCNs allowed us to examine the activity of the genes encoding the NSL complex within different cell type contexts.^40^ Cell type enrichments annotated to modules containing NSL genes when present were almost always neuronal. These findings support *in vitro* experimental results demonstrating changes in multiple markers of mitochondrial dysfunction following NSL KD in neuronal models.^4^ Differences in activity in non-neuronal cell types needs to be validated further, perhaps by examining single-cell RNA-sequencing datasets. Published bulk RNA-sequencing data from CRISPRi-mediated *KAT8* KD in THP-1 cells – a macrophage-like cell line – indeed showed no significant change in *PINK1* expression, contrary to the aforementioned decrease detected in neuroblastoma cells.^6^

GCN analysis was augmented by the use of ARACNe, a method for reverse engineering gene regulatory networks.^63^ By applying ARACNe-AP to members of the NSL complex, we identified significant overlaps between the predicted regulons, increasing our confidence in their representativeness of NSL complex function. Importantly, the NSL regulon, comprising genes predicted to be regulated by three or more NSL genes, was enriched for multiple pathways genetically linked to Parkinson’s disease through common genetic variation and almost enriched for heritability through stratified LDSC.^64^ This strengthens the notion that the NSL complex acts across the genome to regulate the expression of multiple different genes linked to Parkinson’s disease risk. eQTL analysis may be a way of testing this result more directly, by asking whether SNPs regulating the expression of NSL genes in *cis* also regulate the expression of genes relating to Parkinson’s disease in *trans*. However, very large sample numbers are required to power such an analysis.

Individual genes predicted to be regulated by the NSL complex served as prioritised targets for *in vitro* validation. Significant changes were detected in the expression of seven Parkinson’s disease-associated genes following *KANSL1* and *KAT8* KD, suggesting the NSL complex may act as a master regulator of multiple pathways implicated in disease pathology. Furthermore, several of these regulatory relationships have been captured in existing literature: RNA-sequencing of THP-1 cells found a significant decrease in *DGKQ* expression following *KAT8* KD, though the changes in NCKIPSD and WDR45 expression were not confirmed.^6^ Similarly, RNA-sequencing of embryonic fibroblasts from *KANSL1* knock-out mice also captured a decrease in *PINK1*, *BIN3* and *DGKQ*, but found contrasting increases in *CSTB* and *NCKIPSD*.^66^ These results support the notion that the gene regulatory activity of the NSL complex has potential organism and cell-type specificity. Parallel assay for transposase-accessible chromatin (ATAC)- or chromatin immuno-precipitation (ChIP)-sequencing experiments would no doubt assist the characterisation of NSL complex activity at specific loci.

In this study, we demonstrate the potential of *in silico* analyses to identify regulatory relationships between genes highlighted in GWASs and successfully validate such interactions *in vitro*. The NSL complex thus provides a potentially useful therapeutic target which could be used to simultaneously modulate a range of molecular pathways implicated in Parkinson’s disease, a strategy which might combat problematic compensatory responses that can occur when individual pathway components are targeted. Chromatin regulation is already an emerging target for the treatment of rare disorders including specific cancers and developmental diseases, and we believe a similar approach may prove effective in neurodegenerative disorders.^67–69^

## Abbreviations

ARACNe-AP: Algorithm for the Reconstruction of Accurate Cellular Networks with adaptive partitioning
ATAC: assay for transposase-accessible chromatin
BP: biological process
CC: cellular component
ChIP: chromatin immuno-precipitation
DMEM: Dulbecco’s Modified Eagle Medium
eQTL: expression quantitative trait locus
EWCE: expression weighted cell-type enrichment
FBS: foetal bovine serum
FDR: false discovery rate
GCN: gene co-expression network
GMSCA: gene multifunctionality in secondary co-expression network analysis
GO: gene ontology
GTEx: Genotype Tissue Expression
GWAS: genome wide association study
iPSC: induced pluripotent stem cell
kb: kilobase
LDSC: linkage disequilibrium score regression
Mb: megabase
ME: module eigengene
MF: molecular function
MSL: male-specific lethal
ns: not significant
NSL: non-specific lethal
NT: non-transfected
pUb: phospho-ubiquitin
SAPE: Streptavidin R-Phycoerythrin conjugate
siRNA: short interfering RNA
SNP: single nucleotide polymorphism
TPM: transcripts per million
WGCNA: weighted gene co-expression network analysis
UKBEC: United Kingdom Brain Expression Consortium

## Supplementary material

Supplementary materials are included in a separate PDF file.

## Author notes

Mina Ryten and Helene Plun-Favreau contributed to this work equally.

---

**Supplementary Figure 1.**
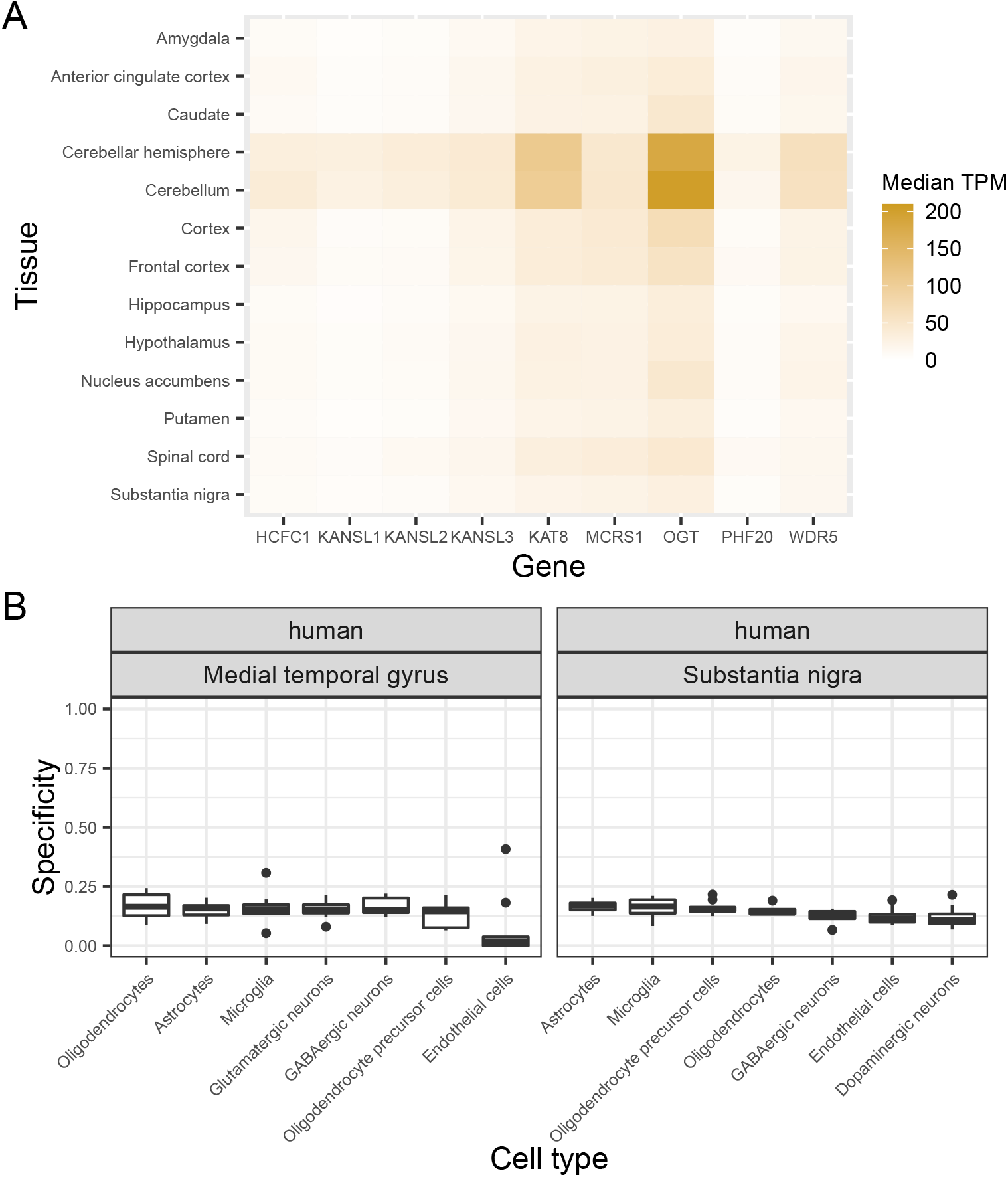
Tissue and cell type specificity of NSL complex gene expression. (A) Median TPM for each NSL gene in brain tissue data obtained from GTEx protal. (B) Plot of specificity values for NSL genes across brain-related cell types in two independent datasets, obtained from medial temporal gyrus and substantia nigra tissue. Data was obtained from Hodge et al., 2019 and Agarwal et al., 2020.^31,32^ Genotype Tissue Expression (GTEx), nonspecific lethal (NSL), transcripts per million (TPM).

**Supplementary Figure 2.**
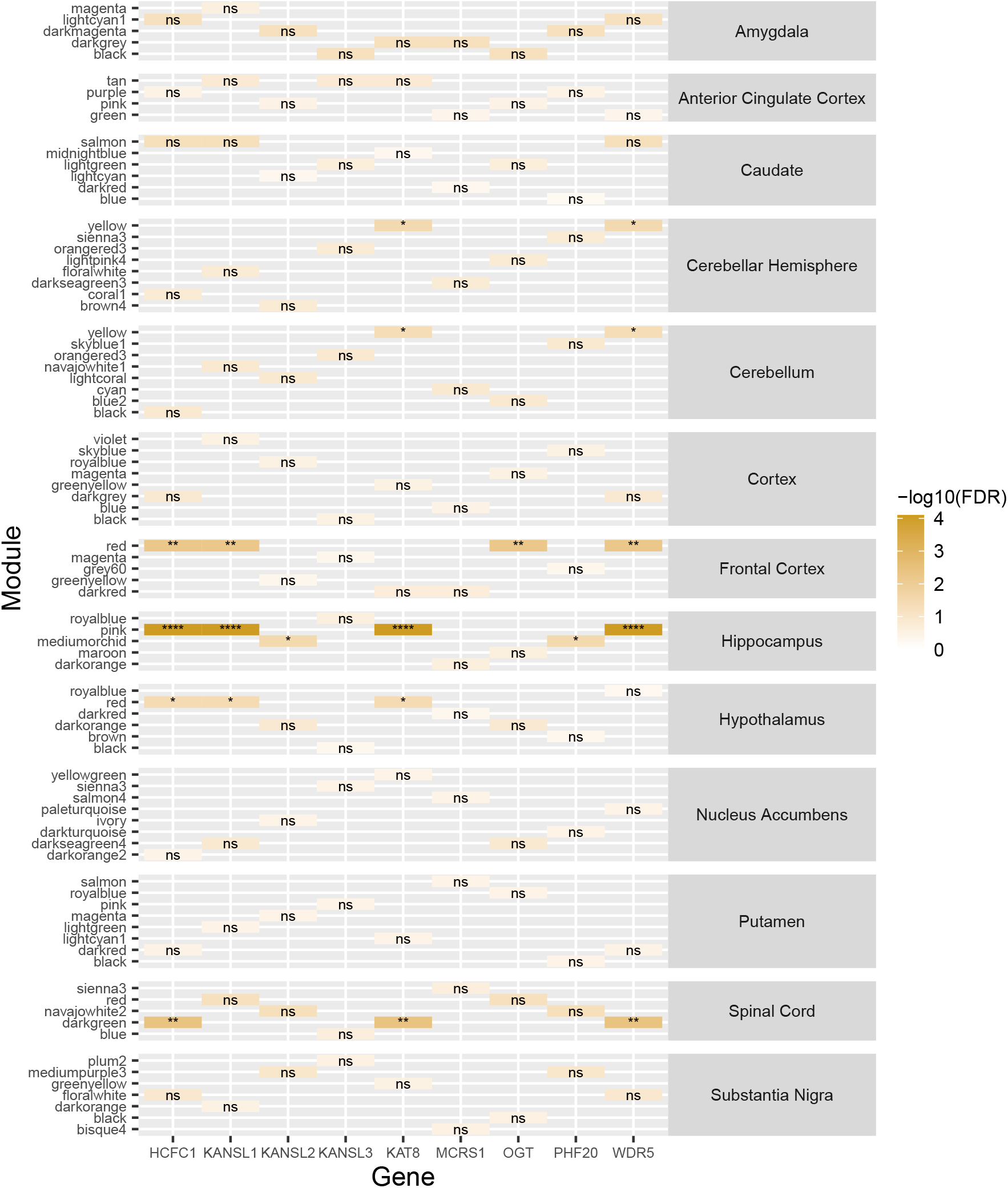
NSL complex gene enrichment across gene co-expression networks generated using GTEx data from multiple CNS regions. GCNs constructed using GTEx datasets containing transcriptomic data from 13 CNS regions. Fishers exact test, displayed over a log scale (FDR-corrected p-values). ns, FDR > 0.05; *, FDR < 0.05; **, FDR < 0.01; ***, FDR < 0.001; ****, FDR < 0.0001. False discovery rate (FDR), gene coexpression network (GCN), Genotype Tissue Expression (GTEx), non-specific lethal (NSL).

**Supplementary Figure 3.**
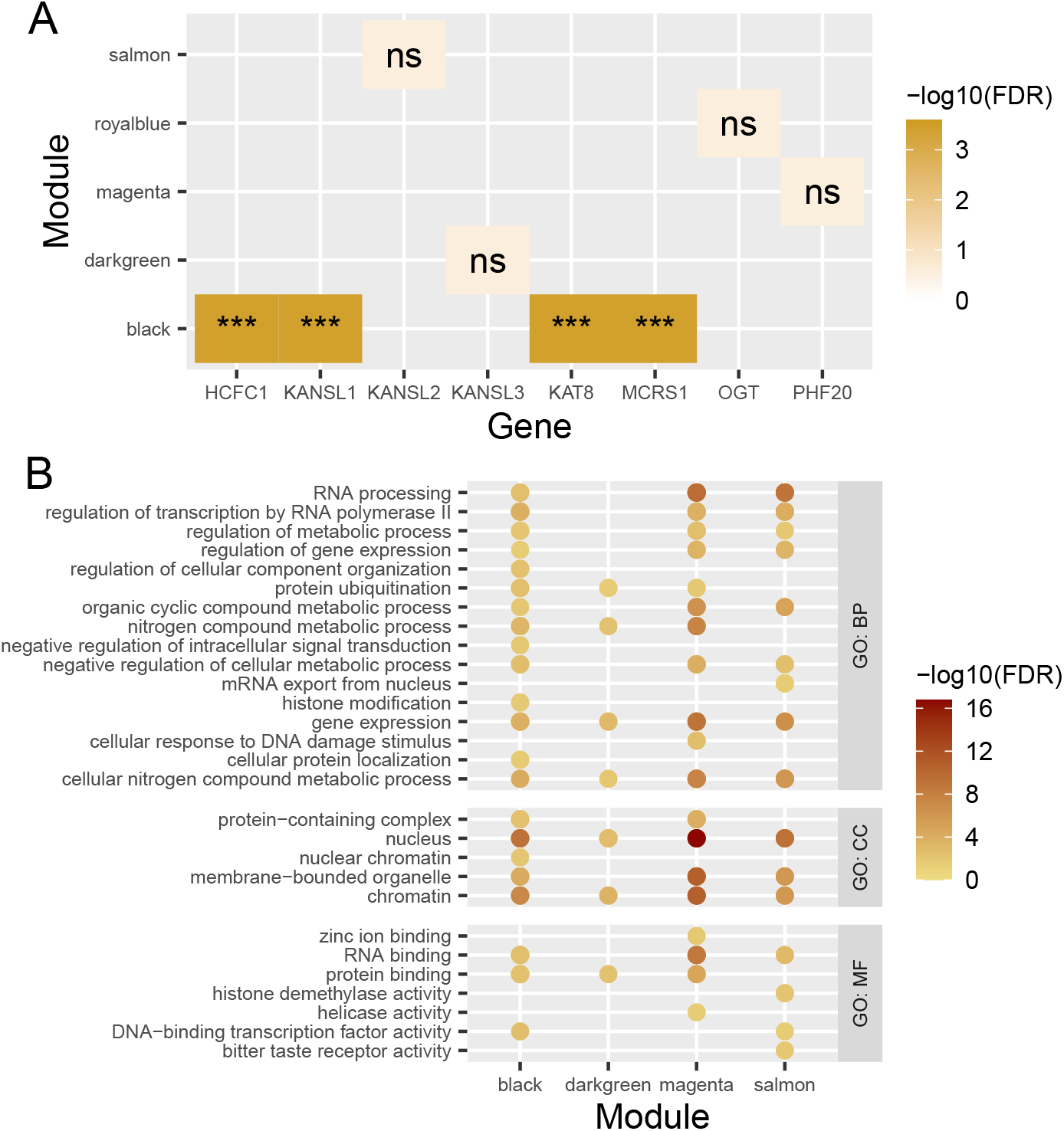
Exploration of UKBEC frontal cortex gene co-expression network. (A) NSL gene enrichment analysis across GCN modules constructed using UKBEC datasets. (B) GO BP, MF and CC term enrichments for GCN modules containing NSL genes. Each term has been uniformly reduced to a parent term in order to group together similar terms. Colour corresponds to the p-value of the most significantly enriched child term within the parent term. Fishers exact test, displayed over a log scale (FDR-corrected p-values). ns, FDR > 0.05; *, FDR < 0.05; **, FDR < 0.01; ***, FDR < 0.001; ****, FDR < 0.0001. Biological process (BP), cellular compartment (CC), false discovery rate (FDR), gene co-expression network (GCN), gene ontology (GO), molecular function (MF), non-specific lethal (NSL), United Kingdom Brain Expression Consortium (UKBEC).

**Supplementary Figure 4.**
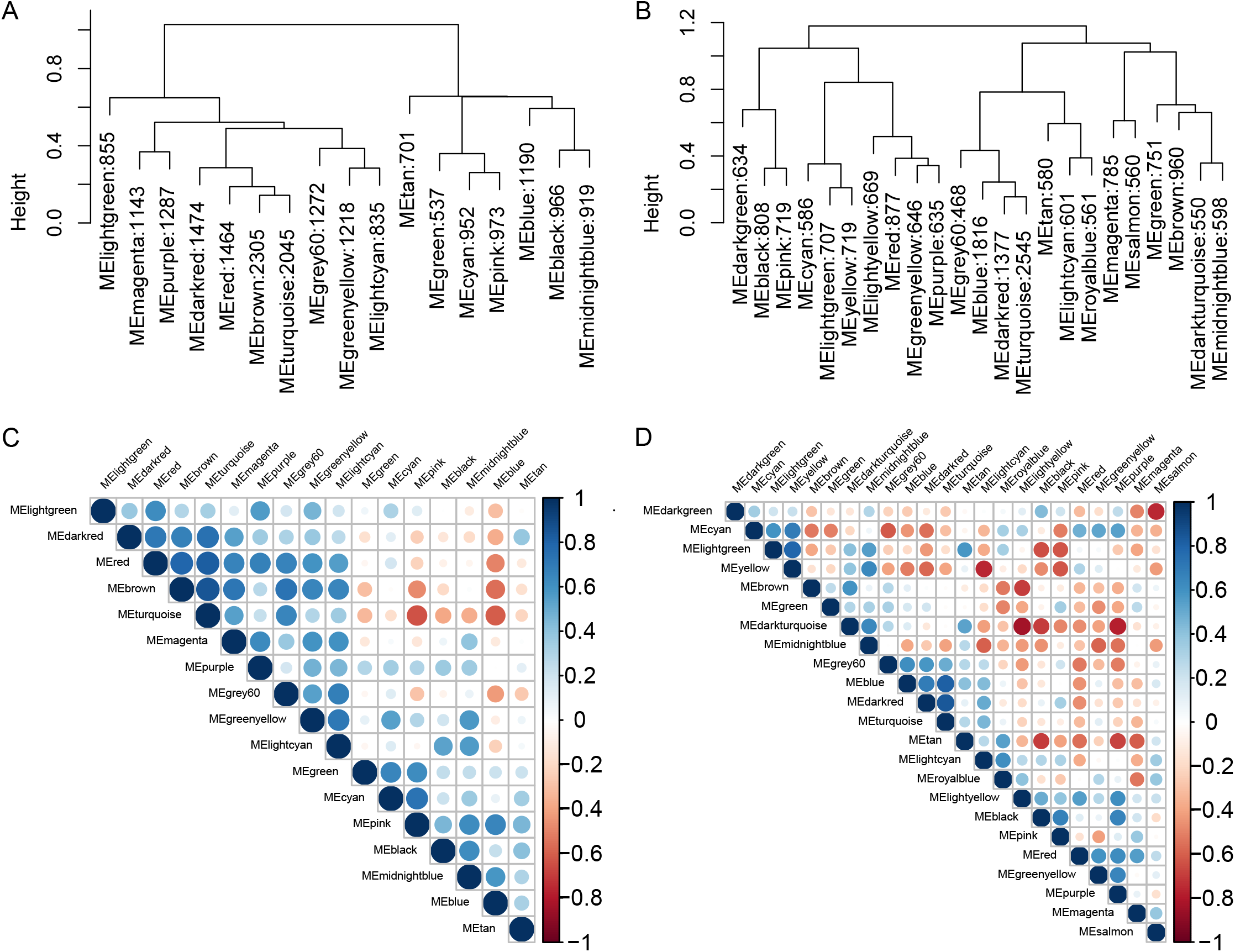
Inter-module relationships within frontal cotrex gene coexpression networks. (A-B) GCN module eigengene dendrogram, labelled as ME module: number of genes within module. (C-D) Spearman’s rank correlation plot for GCN modules, labelled as ME module. Colour corresponds to R number and size of circle corresponds to p-value significance of association. (A,C) describe GTEx GCNs and (B,D) describe UKBEC GCNs. Gene co-expression network (GCN), Genotype Tissue Expression (GTEx), module eigengene (ME), United Kingdom Brain Expression Consortium (UKBEC).

**Supplementary Figure 5.**
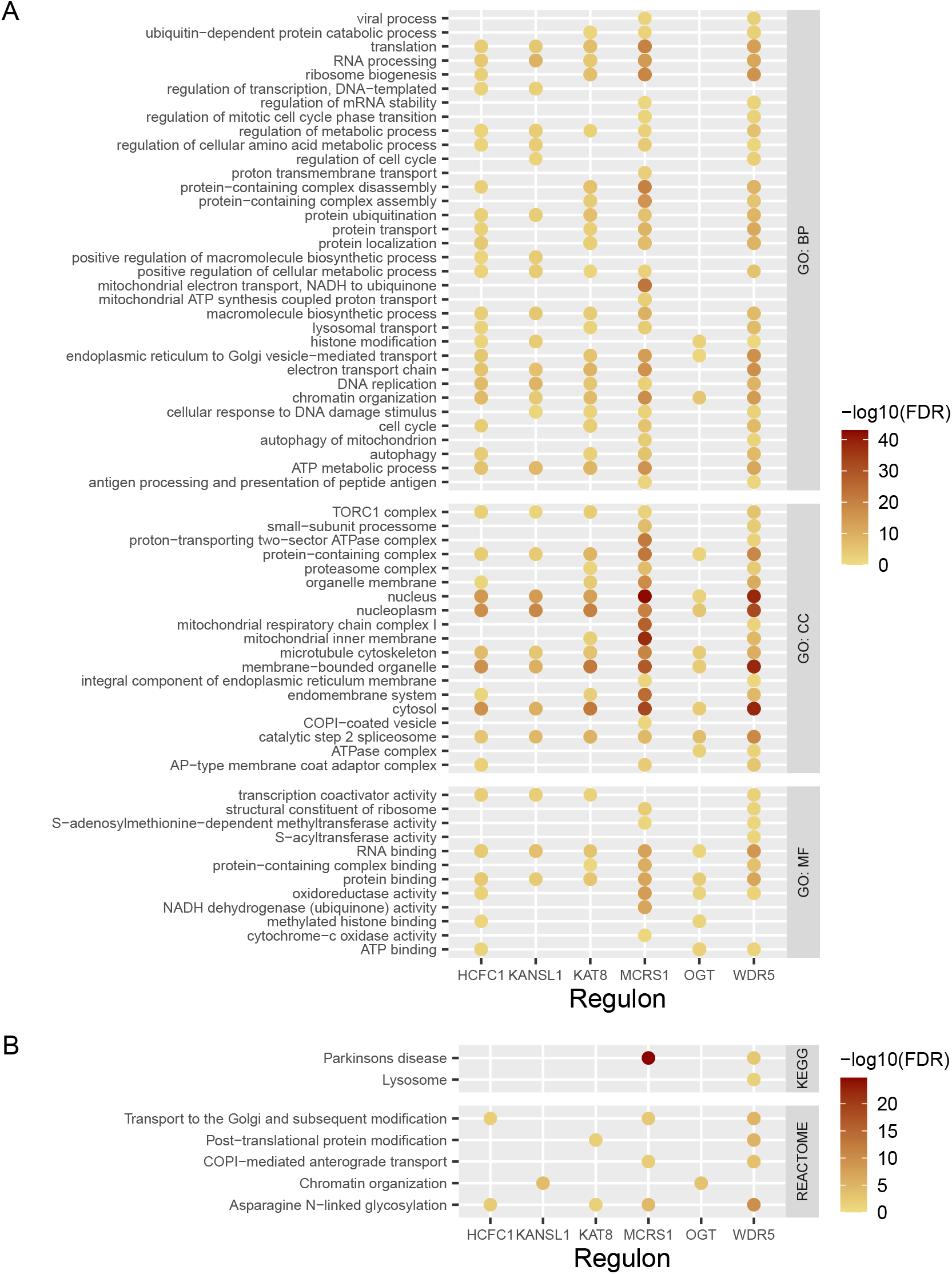
Gene set enrichment analysis of individual regulons of NSL genes, derived from ARACNe-AP analysis of ‘red’ and ‘darkred’ GTEx primary gene coexpression network modules. (A) GO BP, MF and CC term enrichments for each NSL regulon. Each term has been uniformly reduced to a parent term in order to group together similar terms. Colour corresponds to the p-value of the most significantly enriched child term within the parent term. (B) REACTOME and KEGG term enrichments for each NSL regulon, filtered for those genetically linked to Parkinson’s disease. Fishers exact tests, displayed over a log scale (FDR corrected p-values). ns, FDR > 0.05; *, FDR < 0.05; **, FDR < 0.01; ***, FDR < 0.001; ****, FDR < 0.0001. Biological process (BP), cellular compartment (CC), false discovery rate (FDR), gene co-expression network (GCN), gene ontology (GO), Genotype Tissue Expression (GTEx), molecular function (MF), non-specific lethal (NSL).

**Supplementary Figure 6.**
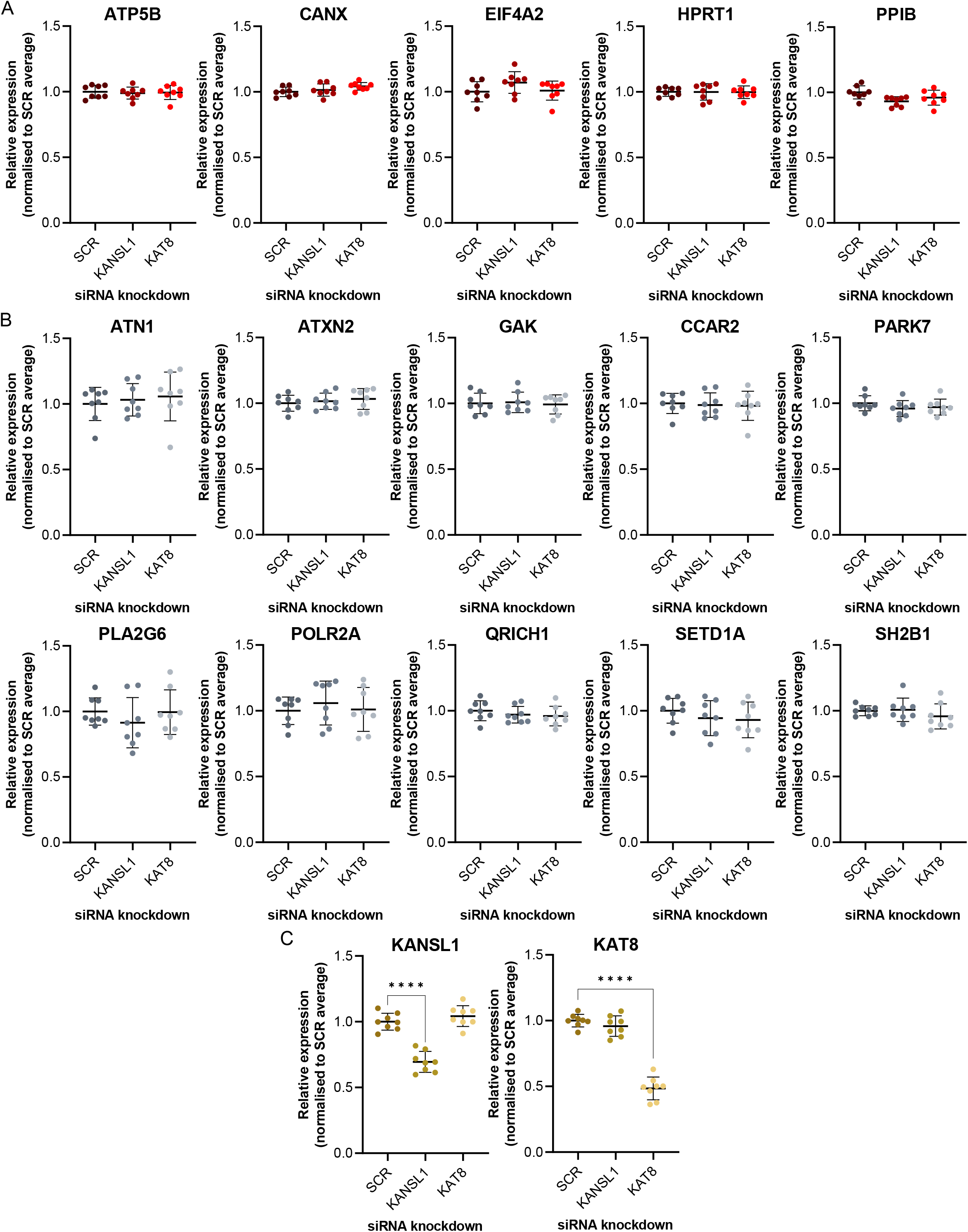
Changes in mRNA levels of housekeeping, NSL complex and Parkinson’s disease-associated genes following knockdown of NSL complex genes. (A) Housekeeping genes, (B) Parkinson’s disease-associated genes, (C) NSL genes. Measured using QuantiGene multiplex assay, normalised to average SCR control. One-way ANOVA with Dunnett correction for multiple comparisons, n=8. *, p<0.05; **, p < 0.01, ***, p<0.001; ****, p<0.0001. Non-specific lethal (NSL), scrambled (SCR).

**Supplementary Table 1.** Summary of results from ARACNe-AP analysis of GTEx gene co-expression network modules. Total number of targets predicted to be directly regulated by each NSL gene. Genotype Tissue Expression (GTEx), non-specific lethal (NSL).

| Regulon | Number of genes |
| --- | --- |
| HCFC1 | 704 |
| KANSL1 | 491 |
| OGT | 617 |
| KAT8 | 775 |
| MCRS1 | 1584 |
| WDR5 | 1788 |

**Supplementary Table 2.** Enrichment of Parkinson’s disease SNP-based heritability within cumulative frequencies of NSL complex regulons derived from ARACNe-AP analysis of GTEx data. Stratified LDSC analysis generated enrichment values, representing the proportion of heritability accounted for by the annotation after controlling for all other categories in the model. These were used to calculate two-tailed p-values, which were FDR-corrected. False discovery rate (FDR), Genotype Tissue Expression (GTEx), linkage disequelibrium score regression (LDSC), non-specific lethal (NSL), single nucleotide polymorphism (SNP).

| Regulon cumulative frequency | Enrichment |  | Regression coefficient |  |
| --- | --- | --- | --- | --- |
|  | P-value | FDR | P-value | FDR |
| 1 or more | 2.65x10<sup>-4</sup> | 5.32x10<sup>-4</sup> | 0.402 | 0.803 |
| 2 or more | 2.16x10<sup>-4</sup> | 5.32x10<sup>-4</sup> | 0.317 | 0.803 |
| 3 or more | 2.54x10<sup>-4</sup> | 5.32x10<sup>-4</sup> | 0.0519 | 0.312 |
| 4 or more | 0.0314 | 0.0471 | 0.768 | 0.921 |
| 5 or more | 0.113 | 0.136 | 0.968 | 0.968 |
| 6 | 0.268 | 0.268 | 0.619 | 0.921 |

**Supplementary Table 3.** Enrichment of Parkinson’s disease SNP-based heritability within individual NSL complex regulons derived from ARACNe-AP analysis of GTEx data. Stratified LDSC analysis generated enrichment values, representing the proportion of heritability accounted for by the annotation, and regression coefficients, representing the contribution of an annotation after controlling for all other categories in the model. These were used to calculate a two-tailed p-values, which were FDR-corrected. False discovery rate (FDR), Genotype Tissue Expression (GTEx), linkage disequilibrium score regression (LDSC), single nucleotide polymorphism (SNP).

| Regulon | Enrichment |  | Regression coefficient |  |
| --- | --- | --- | --- | --- |
|  | P-value | FDR | P-value | FDR |
| HCFC1 | 5.11x10<sup>-4</sup> | 1.53x10<sup>-3</sup> | 0.0239 | 0.143 |
| KANSL1 | 4.03x10<sup>-3</sup> | 6.38x10<sup>-3</sup> | 0.270 | 0.540 |
| KAT8 | 0.0118 | 0.0118 | 0.862 | 0.862 |
| MCRS1 | 5.32x10<sup>-3</sup> | 6.38x10<sup>-3</sup> | 0.808 | 0.862 |
| OGT | 4.41x10<sup>-3</sup> | 6.38x10<sup>-3</sup> | 0.232 | 0.540 |
| WDR5 | 2.82x10<sup>-4</sup> | 1.53x10<sup>-3</sup> | 0.463 | 0.695 |

